# Logging disrupts the ecology of molecules in headwater streams

**DOI:** 10.1101/2023.03.07.531469

**Authors:** Erika C Freeman, Erik JS Emilson, Kara Webster, Thorsten Dittmar, Andrew J Tanentzap

## Abstract

Global demand for wood products is increasing forest harvest. One understudied consequence of logging is that it accelerates mobilization of dissolved organic matter (DOM) from soils to aquatic ecosystems. Here, we tested how logging changed DOM in headwaters of hardwood-dominated catchments in northern Ontario, Canada. We apply a before-after control-impact experiment across four catchments for three years. DOM concentration in streams from logged catchments quadrupled, on average, after the first year post-harvest, but resulting changes to the molecular composition of DOM persisted for at least two-years. Ultrahigh-resolution mass spectrometry revealed that DOM within logged catchments was more energy-rich and chemically diverse than in controls, with novel highly unsaturated polyphenols, carboxylic-rich alicyclic, and nitrogen-containing formulae. The molecular composition of stream DOM measured fortnightly post-harvest was most strongly associated with DOM composition within intermediate and deeper layers of contributing soils, likely due to increased hydrological connectivity post-harvest. We estimate logging increased the total annual flux of dissolved organic carbon in streams by 6.4% of extracted wood carbon, and this carbon was more likely to be released into the atmosphere. Carbon accounting of forestry, including as a natural climate solution, must now consider the transport and fate of DOM from land into water.

## Main

Economists predict that the demand for wood products will increase by 19% in the coming decade^1, 2^, especially with the shift towards using harvested wood products to store carbon as a natural climate solution^3, 4^. Harvest of wood from plantations and secondary and primary forests must rise to meet this demand^2, 5^, but may have unintended consequences for carbon sequestration. Forest removal and soil disturbance associated with wood harvest typically increase dissolved organic matter (DOM) export to aquatic ecosystems^6, 7^, though decreases have been reported^8, 9^, especially in tropical systems^10^. Even without considering harvest-associated increases in DOM export, at least 15% of annual terrestrial net ecosystem productivity is exported from soils into inland waters, which is conservative, as this estimate only considers mineral soils^11^. Once in waters, DOM is highly reactive^12, 13^. Compared with soils, aquatic DOM is more likely to be respired rather than used for microbial growth^14^, making terrestrial carbon more susceptible to re-release to the atmosphere^13,14,15^. Therefore, the quantity and composition of DOM exported from soils into waters after logging may offset the intended benefits of natural climate solutions alongside changing how aquatic ecosystems function.

How logging changes DOM export and composition is largely unknown because DOM is a highly complex mixture. Each compound in a DOM mixture has unique intrinsic properties and interacts differently with physicochemical and biological processes^16^. For these reasons, each compound has a different likelihood of being degraded (i.e., assimilated into microbial biomass or respired and potentially re-released to the atmosphere as CO_2_ or CH_4_), flocculating into larger mixtures, or deposited into sediment^17^. Advances in ultra-high-resolution mass spectroscopy (UHR-MS) now provide an opportunity to characterise the potential fate of compounds within DOM after logging. However, the fate of each compound will depend on other compounds present in a mixture of DOM^16^. These dependencies among compounds can be explored under the framework of the “ecology of molecules”^16^. This framework predicts that the reactivity and eventual fate of DOM is an emergent property of the interactions among compounds, such as measured by the presence of unique molecular formulae in a mixture (termed “chemodiversity”) and the proportion of different compound classes, typically identified based on elemental ratios of individual formulae^18^. Previous studies suggest that both chemodiversity and compound composition alter microbial and physicochemical reactivity^18, 19^ and predict carbon emissions and the productivity of aquatic food webs^20–22^. For example, carboxylic-rich alicyclic molecules (CRAMs) containing relatively intermediate H:C and O:C ratios tend to be less reactive, and so may persist longer in the environment^23, 24^, whereas high H:C (>1.5) aliphatic, protein-like, lipid-like, and high N and P-containing formulae^25–27^ are preferentially degraded by microbes and so may be more easily respired^28, 29^. By interpreting UHR-MS data within the ecology of molecules framework, we can begin to identify how logging changes the molecular properties of DOM and the implications for forest carbon budgets and functioning of aquatic ecosystems.

Disturbance that mobilises terrestrial organic matter into downstream waters risks introducing otherwise stable carbon into the pool of reactive DOM and increasing the likelihood of carbon being released to the atmosphere. Historically, aquatic DOM sourced from terrestrial ecosystems was assumed to be resistant to biodegradation and therefore escape atmospheric release^30, 31^. This view has been revised by evidence that microbes can rapidly oxidise terrestrial DOM once it is released into aquatic ecosystems, even when it is thousands of years old^32–34^. Previous research has mostly studied how forest harvest changes DOM fluxes, such as by exporting organic matter from deeper soils^35–37^, with few studies investigating changes in DOM quality, mostly though fluorescence characteristics. The studies suggest either no detectable change in DOM post-harvest^38^, or an increase in protein-like compounds attributed to less woody debris post-harvest^15, 39^. However, fluorescence data provide little information on DOM diversity and do not disambiguate sources within terrestrial systems. The application of UHR-MS offers a major advance on these approaches by helping to identify the mechanisms that contribute to changes in DOM composition. Changed DOM composition in streams after forest harvest could indicate greater contribution by surface flowpaths, e.g. overland flow and near-surface lateral flow^40, 41^. Increased precipitation entering the soil following tree removal also enhances soil wetness and the expansion of surface saturated areas^42^. Further alteration in DOM sources can be expected due to soil disturbance, changes in nutrient dynamics, and/or inputs of woody debris leftover by harvest activities^43^. Knowledge of these mechanisms are essential to design forest management practices that protect aquatic ecosystems and minimize the potential of downstream carbon emissions.

First-order streams are ideal to begin investigating the mechanisms driving DOM composition after land-use change because they carry a stronger imprint of their surrounding watersheds than higher order waterways^44, 45^ and their terrestrial inputs are simpler to resolve. First-order streams are also important for carbon cycling, making up ca. 36% of the carbon emissions from running waters^46^. Forest harvest is well known to increase DOM export^7^ in first-order streams by modulating the factors controlling soil solution concentrations of DOM^47, 48^, such as temperature^53^, moisture^49^, logging residues^48^, and by increasing lateral groundwater flow from deeper to more superficial soils rich in DOM^50–52^, particularly in temperate hardwood forests^6, 53–55^. The same processes may impact DOM composition and chemodiversity. For example, if hydrology changes, so too should the soil layers that water interacts with before entering streams. Soil DOM can become homogenised along soil depth and hillslope position gradients^43^, and so should respond similarly to water that interacts with deeper soil layers after harvest. In river networks, Lynch at al.^56^ also observed a loss in DOM chemodiversity with increasing flow conditions because of reduced physical opportunities for microbial metabolism. Given the analogous increase in flow under post-harvest conditions, logging could produce a similar homogenizing effect on DOM composition, i.e. lower chemodiversity.

Here, we experimentally tested how DOM composition and chemodiversity changes after forest harvest in first-order streams and surrounding hillslope soils over three years. We replicated our study in four hardwood-dominated catchments, two of which were selectively harvested with paired unharvested controls selected for nearly identical climate, underlying geology, size, and topography (see Methods). We quantified spatial and temporal differences in DOM molecular composition before and after forest harvest using a before-after-control-impact statistical design and complementary analytical chemistry approaches, namely Fourier-transform ion cyclotron resonance mass spectrometry (FT-ICR MS) and fluorescence spectroscopy. We report large changes in DOM composition after forest harvest and their likely sources via changes to soil flow paths, thereby identifying practices for forest management aimed at climate change mitigation, such as with nature-based solutions^32–34^.

### The immediate mobilisation of DOM after forest harvest

Consistent with the expectation^7^ that forest harvest increases DOM export to streams, we found DOM concentrations were elevated in the logged catchments relative to the controls in the year of harvest (Figure 1, Table S1). This selective forest harvest (see Methods) largely offset the decline in DOM seen in the controls after increased seasonal discharge^50, 51^ (Figure 1b). The rate of change in concentrations observed in the harvested streams (0.08 mg C L^-1^ d^-1^; SE: ± 0.31 mg C L^-1^ d^-1^) was large enough that the mean DOM concentration increased by 100% from initial values (1.22 ± 0.38 mg C L^-1^ d^-1^) and by 600% relative to the controls by the end of the growing season (Figure 1b). Cation and heavy metal concentrations increased similarly in the streams of the harvest sites, indicative of increased nutrient leaching from disturbed soils^6^ (Table S2).

**Figure 1:**
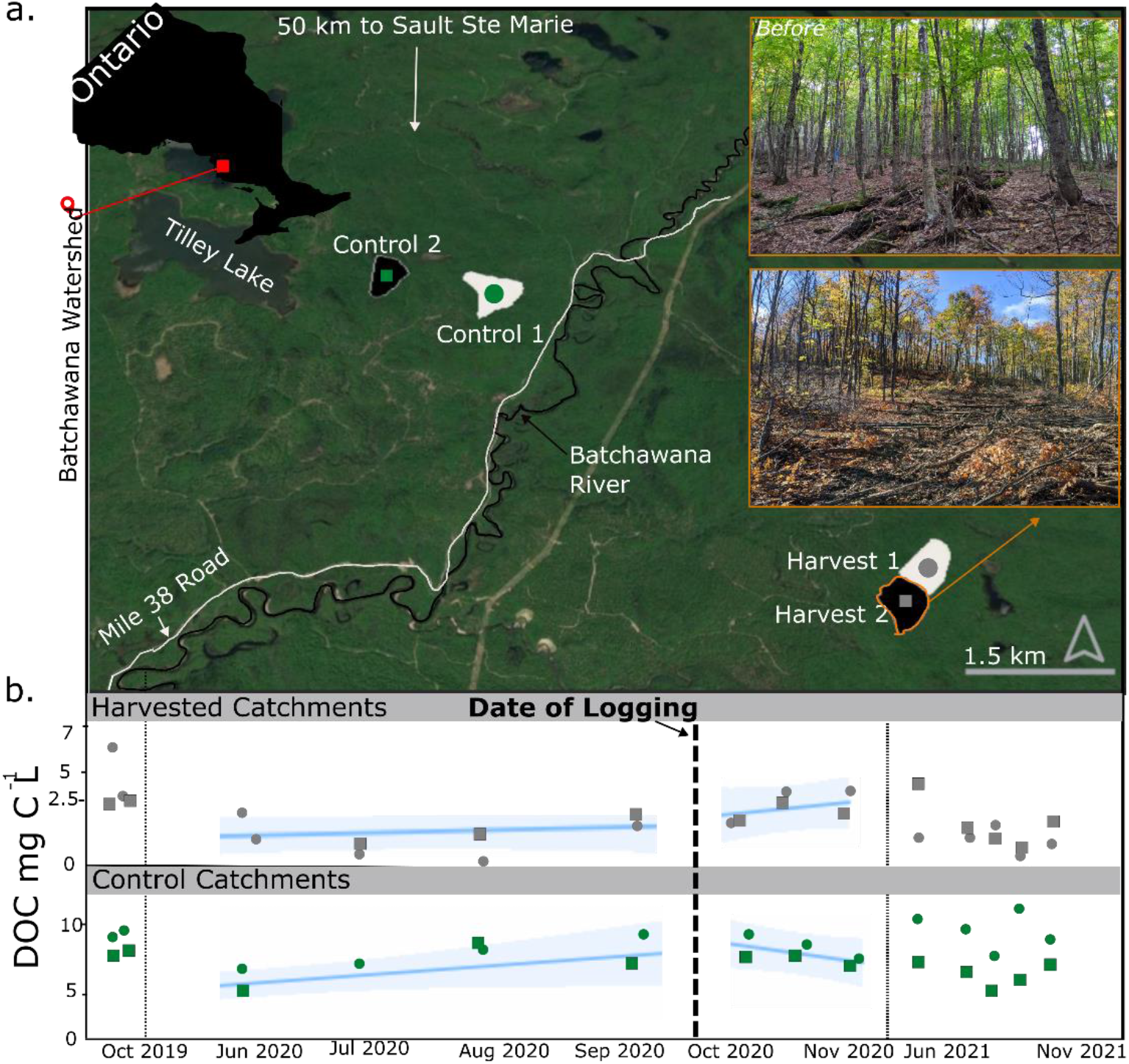
DOM concentrations were elevated in streams of the harvest relative to the control sites immediately after logging. **a.** Map of experimental catchments within the Batchewana Watershed, Ontario, Canada. The four replicate catchments were named based on the impact classification. Control sites were unharvested, and Harvest sites were logged in late September 2020 (bolded dashed line in bottom panel). **b.** Dissolved organic matter (DOM) concentrations faceted by year (2019, 2020, 2021) and treatment (Harvest upper, Control lower). Estimated mean ± 95% confidence interval for temporal trends were plotted as blue lines ± bands where there were statistically significant differences between treatments (Table S1).

To compare the stream DOM fluxes with estimates of carbon removed as wood biomass by logging, we estimated annual carbon export in runoff for the harvested catchments. We multiplied the modelled DOM increase by monthly runoff measured in 9 similarly-sized local catchments (see Methods). We estimated an additional mean 13 kg C month^-1^ ha^-1^ (SE: ± 4 kg C month^-1^ ha^-1^) of DOM was exported from the logged sites compared to the non-harvested catchments. If this increase remained constant for all non-winter months (n=8), an additional ca. 6.4% of the carbon that was harvested as above-ground woody biomass would be lost to the streams solely through increased DOM export (see Methods). We did not account for potential C changes within the soil pool, though previous studies suggest that harvesting reduces soil C by a mean 11.2% (SE: ± 5.7%, n=945)^57^.

### Changes to DOM composition persist after forest harvest

Our results suggest that while there is an increase in DOM concentration only in the year of harvest, the molecular composition of DOM in streams remained influenced by forest harvest well after the initial DOM pulse (Figure 1b; Table S3). The humification index (HIX), which is a unitless measure of the complexity of DOM, increased by 22% from a mean of 4.58 (SE: 1.01) before the harvest to 5.58 (SE: 0.92) within 2 months after harvest. This result indicated that DOM was becoming more polycondensed (lower H:C ratio), or, more soil-like^58^. The increase in HIX was a mean 3.80 (SE: 1.75) larger than in the controls between 2019 to 2021. In contrast to the increase post-harvest, HIX declined between 2019-2021 by a mean of 17% in the control (SE: 1.2) from a mean 16.05 (SE: 1.27) to a mean 13.24 (SE: 0.92), indicating long-term leaching of soil-DOM only into the harvested streams.

Using UHR-MS of solid-phase extractable DOM, we deconstructed the fluorescence signal into individual molecular formulae. We detected 7444 distinct molecular formulae across all streams sampled in 2021. While optical indices reflect only the mean or bulk character of DOM^62^, we identified a specific fraction of the overall molecular formulae (13 and 16%) that were positively and negatively correlated with HIX, respectively (Figure 2a). We then correlated commonly used descriptive traits of individual molecular formulae to the strength of the correlation between the relative intensity of each formulae and the HIX value of the corresponding DOM pool across all sampling times. Molecular formulae that were more positively associated with HIX also had larger values of the formula-based estimate of aromaticity (AI_mod_, ρ = 0.62, p <0.001), which indicates the presence of condensed aromatic structures^63^ and confirms that stream DOM was becoming more like surface soil post-logging, confirmed by leaching studies performed at the site^43^. Studies in lake sediment have found that AI_mod_ strongly correlates with the relative abundance of lignin-like compounds^59^, further supporting a terrestrial origin to the HIX signal rather than changes produced in-stream^60, 61^. The association with AI_mod_ was stronger than with any other metric (Figure 2b).

**Figure 2.**
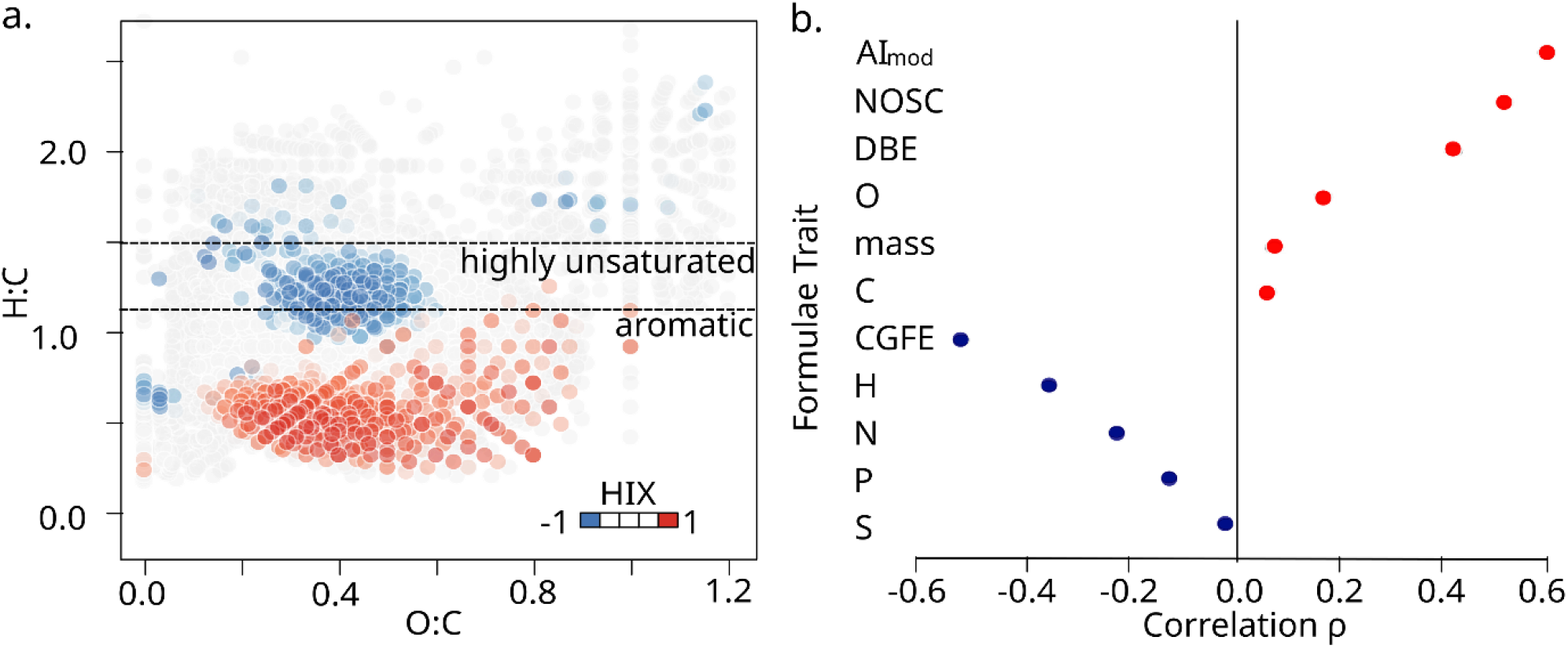
Forest harvest led to persistent increases in the humification index (HIX) of DOM in headwater streams. **a**. Elemental ratios of all molecular formulae detected in our experiment (n=7444). Red and blue points are formulae whose relative peak intensities correlate (p<0.05) positively and negatively with HIX, respectively, based on Spearman rank correlations. Dotted lines indicate aromatic compounds (H:C< 1.1) and the “highly unsaturated” region (1.1 < H:C < 1.5) according to ref^54^. **b**. Spearman rank correlation (ρ) between either the modified aromaticity index (AI_mod_), nominal oxidation state of carbon (NOSC), double bond equivalent (DBE), number of oxygen atoms (O), molecular mass, number of carbon (C) atoms, Gibb’s free energy of carbon oxidation (CGFE), and the number of hydrogen (H), nitrogen (N), phosphorus (P) and sulphur (S) atoms of each molecular formula against the correlation between the relative abundance of each molecular formulae and the HIX of the corresponding DOM pool across all time points (n=7) measured with UHR-MS. For all correlations, p < 0.05.

While the origin of the HIX signal can be inferred, the relative likelihood of logging-added compounds returning to the atmosphere (i.e., its fate) is less clear. Vascular plant-derived compounds that produce strong HIX signals have been reported to have low reactivity^62–64^, but other studies suggest high decay rates within this group^28, 65, 66^. This variation may partly reflect differences in the traits of this compound pool. For example, aromatic compounds (H:C < 1.1) may be more reactive than those in the “highly unsaturated” region (1.1 < H:C < 1.5)^66^. Additionally, the uptake rate of DOM by microbes has been shown in a decomposition experiment to decrease with increasing AI_mod_, but rise when AI_mod_ is between 0.25 and 0.33^66^. While further research is required to understand how reactivity maps to the re-release of carbon to the atmosphere in streams, most of the compounds that correlated with HIX in our study had H:C < 1.1 (Figure 2a) and AI_mod_ > 0.25 (Figure S1), suggesting that the compounds added to streams post-logging may be more reactive than those in pre-logging conditions.

### Increased diversity of harvest-impacted stream DOM

We expected that forest harvest would reduce chemodiversity in streams as DOM became homogenised by high soil hydrological connectivity induced by harvest^56^. However, we found a 32% increase in the number of unique compounds after forest harvest compared to the controls (Figure 3a). The number of unique compounds increased from a mean of 4771 (SE: 330 compounds) to 6292 (SE: 390 compounds), whereas compounds declined in controls from a mean of 5387 (SE: 399 compounds) to 4153 (SE: 260 compounds). We found further support for this diversification post-logging using a multivariate analysis of molecular composition. We calculated the Bray-Curtis dissimilarity between control and harvest sites at each time point. The pairwise distance between paired sites increased after harvest (Figure 3b), suggesting that the molecular composition of DOM was increasingly differentiated. Furthermore, the harvest sites differed more between the before and after samples than the controls (Figure 3). These results are the first direct report that DOM becomes more diverse after forest harvest, and contrast our expectations based on observations from stream networks^56^ that the molecular composition of DOM should become homogenised.

**Figure 3:**
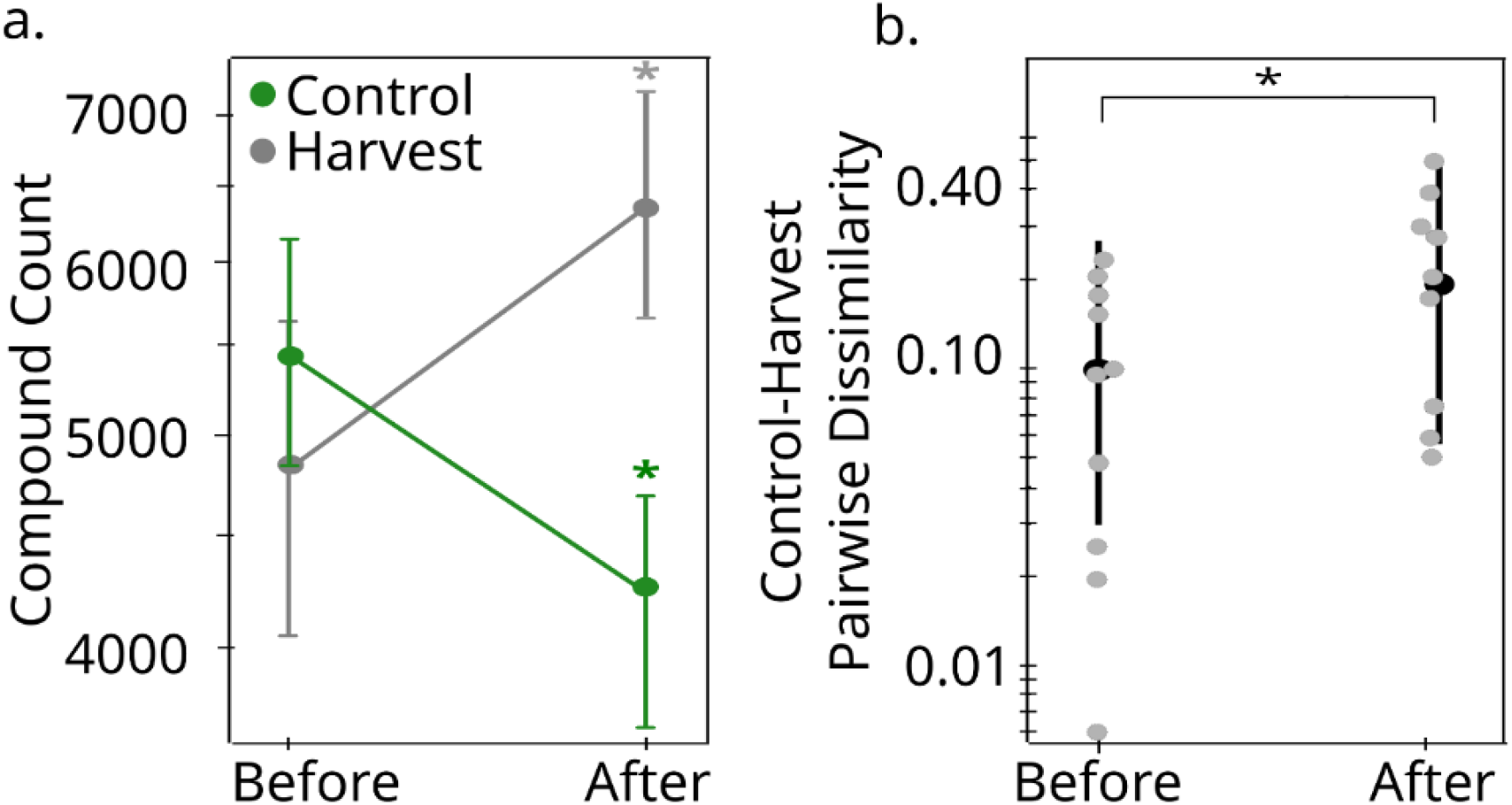
Forest harvest increases molecular diversity of DOM in headwater streams. **a**. Estimated mean (± 95% CI) compound counts in streams draining control and harvested forests before and after logging. The green and grey asterisks indicate statistically significant differences (p<0.05) between control and harvest after logging, and before and after forest logging in both control and harvest sites (Table S4). **b**. Estimated mean (± 95% CI) pairwise dissimilarity between paired control and harvest sites both before and after logging (Table S5). Grey points are individual observations at each time point based on the Bray–Curtis dissimilarity (Figure S2). The asterisk indicates a statistically significant difference (p<0.05) between the mean pairwise dissimilarity before and after logging using a mixed effects model that considered temporal autocorrelation.

We explored the mechanism underlying the unexpected increase in chemodiversity by analyzing the molecular and elemental composition of compounds that were gained in the logged sites after forest harvest. Generally, vascular plant-derived DOM is aromatic and has a low H:C and high O:C ratio^67, 68^. Consistent with this interpretation, we found that the compounds gained after logging had higher O:C ratios (Figure 4a,c). There was also a greater proportion of highly unsaturated phenolic oxygen-rich compounds, i.e., compounds enriched in aromatic structures (Figure 4a), as expected if there was a greater contribution of surficial soil layers and harvest residues to the DOM pool. We also found evidence for contributions from deeper soil layers. On average, N-containing DOM from the logged catchments in this study comprised 43% of the newly added compounds, but only 32% of all identified compounds (Figure 4b, Tables S6, S7). Most of the added N-compounds fell outside of the H:C range interpreted as being associated with autochthonous inputs from primary production^66^ (Figure 4a). Instead, higher N content likely corresponded with release of aged organic carbon^20^ through disturbance of deeper soils. The concomitant increase in carboxyl-rich alicyclic molecules (CRAMs) supports this hypothesis, as the relative intensity of CRAMs increases with microbial processing of carbon and is known to be greater in deeper soils^43, 69^ (Table S7). Together, the input of highly unsaturated phenolic oxygen-rich compounds, N-containing formulae, and CRAMs suggest logging increases chemodiversity downstream by disrupting both surficial and deep stores of soil carbon and adding additional plant material to the soil surface from logging operations. This result builds on the surface contribution model^70^, often posed as an explanation for increased DOM concentrations^7,71^ after harvest. Our results demonstrate that alongside these shallower flowpaths, deeper soils also make contributions that are activated by disturbance.

**Figure 4:**
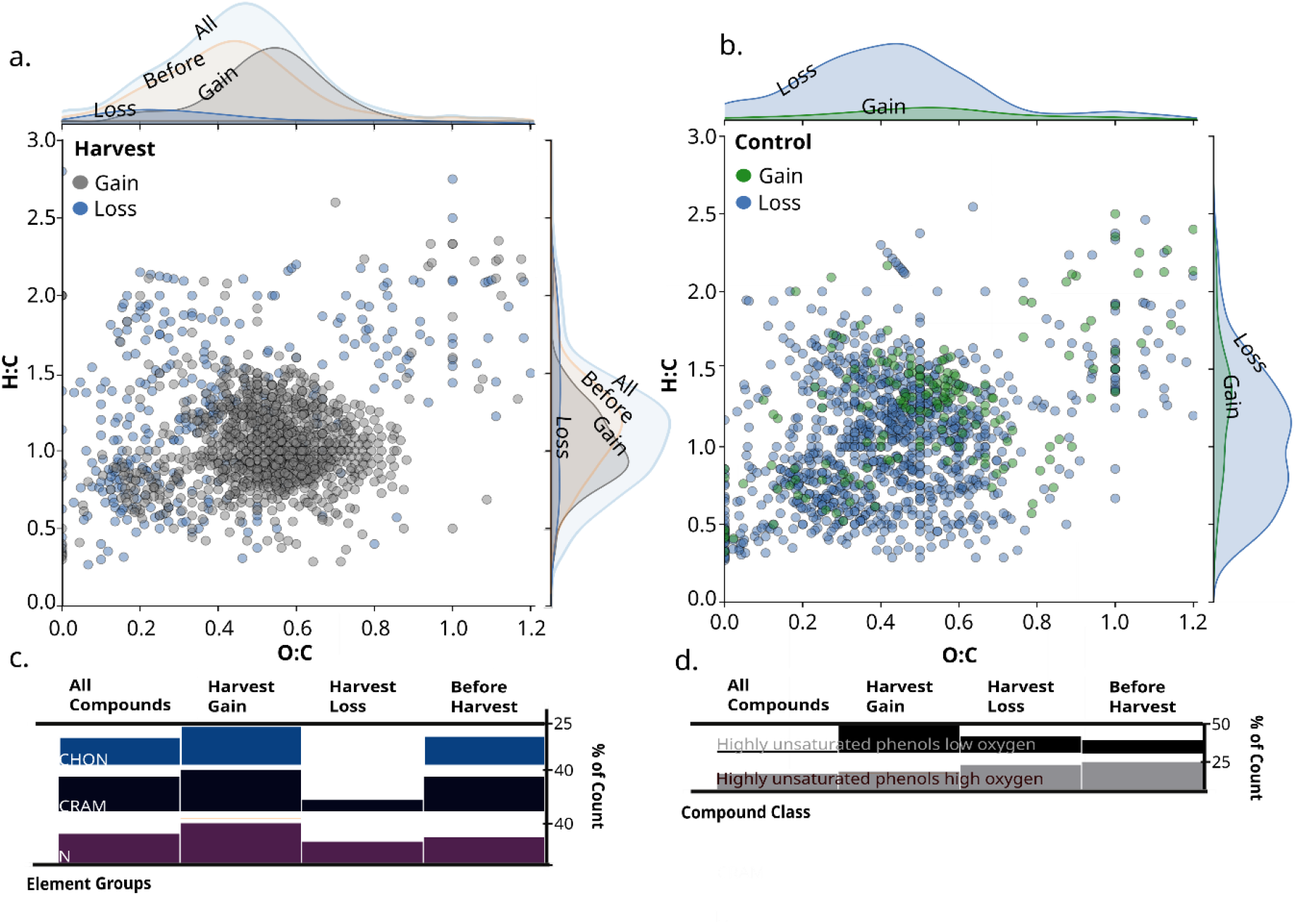
Harvest increases the chemodiversity of stream DOM by introducing compounds reflective of fresh plant material and disturbed soil. Elemental ratios of molecular formulae detected in a. harvest and b. control sites. Probability densities are given along each axis for compounds gained after harvest only in harvest sites (grey, n=1035) and gained after the harvest period only in control sites (green, n=220), for compounds lost only in harvest (blue, n=320) and control (blue, n=962) sites, and all compounds present in the harvest site before logging (n=4927, a. only). We also plotted the percentage of all compounds (n=7444) that contained c. different elements (Table S6) or classified into d. different compound classes (Table S7). Ticks are aligned to the highest bar in each row. The base of the bars equals 0%.

### The terrestrial source of increasing stream chemodiversity

To test further if the increased chemodiversity originated from deeper soils after logging, we tracked DOM from soils into streams monthly during the experiment. We did so by installing lysimeters at 5, 15, 30, and 60 cm depth at shoulder, back, toe, and foot hillslope positions at all four catchments (more details in ref^43^). At each time point, we computed the similarity between all soil samples (n=16) and the stream within a catchment (16 pairs × 4 catchments). We found that the similarity in the molecular composition of DOM between catchment soils and streams persisted in harvest sites compared to controls after logging, suggesting higher hydrological connectivity (Figure 5a). The connectivity between streams and soils in the harvested sites likely arose from intermediate (15 cm) and deep (60 cm) soils remaining connected to streams, given the similarity in DOM composition at these depths in the harvest sites compared to the controls post-logging (Figure 5b). This observation was consistent with the observed increase in nitrogen-rich and CRAM compounds from sub-surface soils in post-harvest conditions (Figure 4). By contrast, streams in unharvested sites declined in their similarity to catchment soils over this same time period from a mean of 60% (95% CI: 58-63%) similarity before logging to 53% (95% CI: 51-56%) afterwards (Figures 5a). The loss in similarity between soils and the stream is consistent with the decline in compound count being attributed to fewer source areas that are hydrologically connected to the control streams (Figure 3a). We observed an increase in compound diversity with harvest (Figure 3a), and infer, based on the similarity in the molecular composition of DOM between streams and soils (Figure 5b), that hydrological connectivity is maintained in harvest sites relative to the controls. The increase in compound diversity with increased hydrological connectivity is the opposite pattern to that observed in rivers, which tended towards less diversity during high-flow periods as more DOM was shunted through the fluvial network^60^.

**Figure 5:**
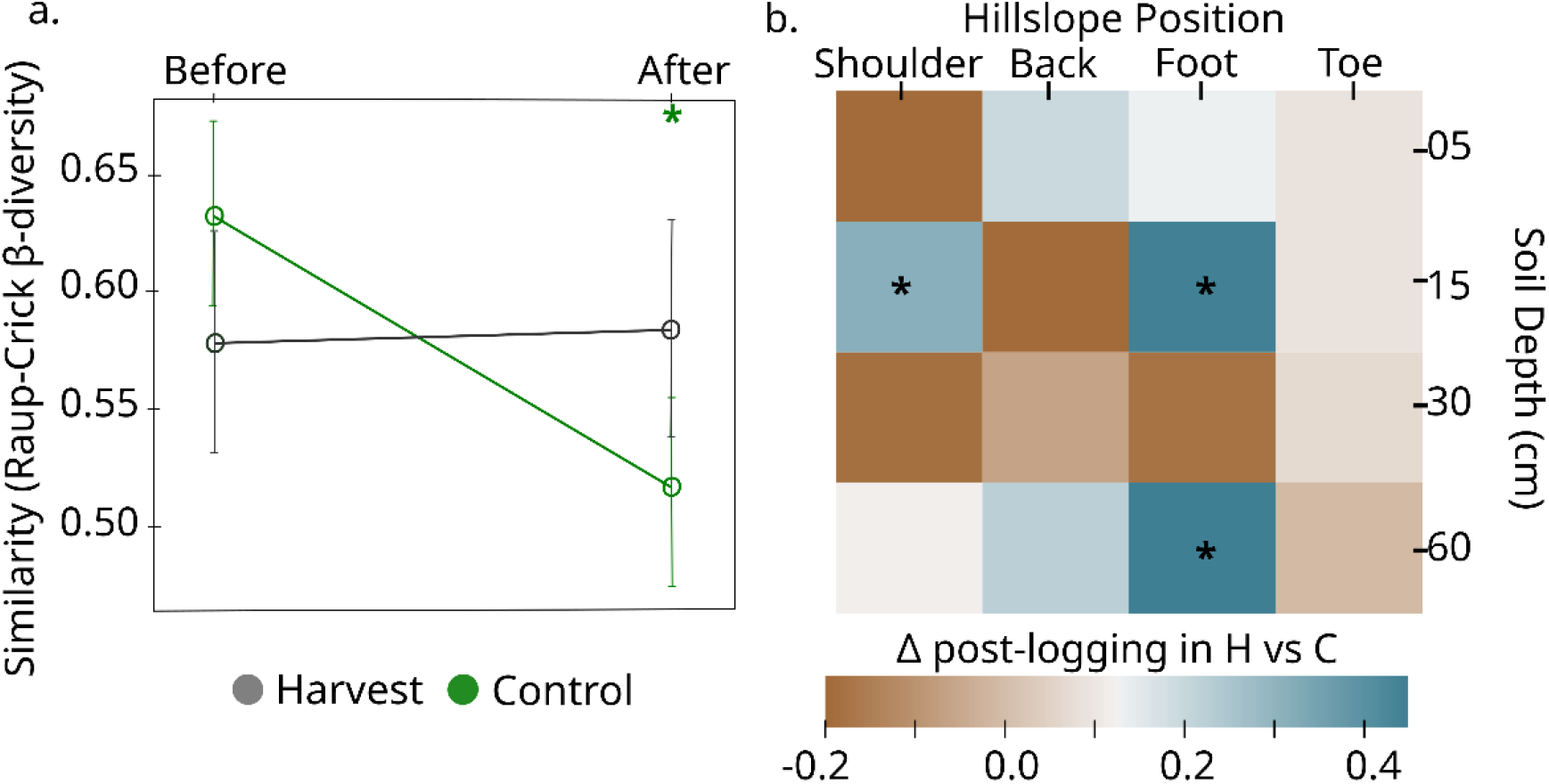
Harvest maintains similarity in DOM composition between stream and soil waters. We calculated the similarity between the molecular composition of DOM in streams and porewater at each soil position (n=4 depths × 4 hillslopes) in each catchment (n=4), accounting for differences solely because of differing numbers of compounds in a sample. **a.** We estimated mean (± 95% CI) similarity across all time points both before (n=5) and after (n=3) logging. * indicates a statistically significant difference (p<0.05) between before and after forest harvest (Table S8). **b**. Changes in molecular composition of DOM after logging in harvest versus the control sites for each depth-by-hillslope combination averaged across catchments. Positive values indicate greater differences in DOM composition after logging in harvest (H) than in control (C) treatments, whereas negative values indicate the reverse. * statistically significant difference in soil-stream similarity between the before and after period between treatments (p<0.05, Table S9).

An alternative explanation for the maintained soil-stream similarity in the harvest sites relative to the controls is that tire or track forces from harvest machinery caused soil displacement and rut formation^78–81^. Rutting may partly be responsible for the signal from deeper soils, as ruts on slopes are preferential routes for runoff, which become deeper because of erosion^72^. By contrast, woody debris left by harvest operations at the soil surface may have contributed to the increased representation of polyphenols^73, 74^. However, only between 15-18% of foliage and fine roots leftover by harvest may be lost annually^75^ and so these sources may have contributed little to DOM changes in our experiment.

### Losing forest soil carbon to the aquatic network

By disrupting soil, forest harvest breaches spatial barriers that would otherwise isolate SOM from aquatic ecosystems. In deeper soil horizons, SOM is typically stabilised via organo-mineral complexation alongside physical disconnection from decomposers and enzymes^76^. Logging activity removes surface soils, exposing deeper layers (as with ruts), or brings them to the surface. The latter outcome results in the breakdown of organo-mineral complexes through the loss of physical protections and the export of deep soil DOM that may be biolabile in aquatic ecosystems^70^. Here we also found that DOM from harvested catchments had unique molecular formulae relative to unharvested catchments and these formulae were absent in the streams prior to harvest. This pulse of unique molecules to streams could have unexpected consequences on their ecology of molecules, analogous to species introductions in biological communities^77^. These unique compounds from typically unconnected soil environments may impact stream microbial communities differently than in the compounds’ native soil “range”, potentially reshuffling the network of relationships amongst DOM compounds and stream microorganisms^19, 78^. Uncovering the relationship between unique (and bioactive) soil-derived additions from deep soils into stream microbial communities is a necessary next step in quantifying aquatic carbon losses to the atmosphere following terrestrial disturbance events such as forest harvest.

Although we did not test the direct consequence of additions of dead organic matter sitting on the soil surface after logging operations (i.e., harvest slash), long-term experiments suggest that much of it may be rapidly leached as DOM in temperate forests^79^. These inputs of leaf litter have also been found, unexpectedly, to prime the microbial community to breakdown additional soil organic matter and change its composition^80, 81^. Leaching of bioavailable plant material through hydrological flow after forest harvest may have similarly unexpected effects on metabolism in water and sediments. For example, Emilson et al^82^ found that fewer dissolved phenolics leaching from terrestrial organic matter increased methane production in aquatic sediments. If inputs of harvest slash reduce soil organic matter through microbial priming, there may also be accompanying changes in the composition of leached DOM that impact aquatic metabolism. Tracking the fate of DOM derived from slash is therefore key to understand the efficacy of natural climate solutions such as forest harvest followed by long-term carbon storage in wood biomass.

### Considerations for forest management practices and policies

Our finding that logging impacts the similarity between soil and stream DOM at a molecular level highlights the need for more integrated land-water management, as well as the need for integration of aquatic fluxes into forest carbon budgets and models. There is a history of managing forests for the protection of water resources in many countries through best management practices (BMPs) primarily targeting the reduction of soil disturbance, the retention of riparian buffer strips, and the reduction of road-related runoff^89^. While these management practices are generally effective at reducing sedimentation^90, 91^, we have demonstrated persistent changes in the molecular composition of DOM, with implications for the fate of soil-derived carbon. Surprisingly, we found that hydrological and biogeochemical alterations occurred within the soil-water interface of logged forests even with low-impact, selection cut techniques in tolerant hardwood stands with BMPs implemented to protect downstream waters. Therefore, our results clearly demonstrate a need to revisit sustainable forest management practices and policies, and forest carbon budget models, to consider leaching of soil carbon into aquatic networks. Without these considerations, the potential carbon sequestration from wood harvest will be miscalculated and have unintended consequences for aquatic ecosystems.

## Methods

### Site description and selection

The four experimental hardwood-dominated catchments were located northeast of Lake Superior, Ontario, Canada (Figure 1) on the northern edge of the Great Lakes-St Lawrence Forest Region^83^ the second largest forest region in Canada. We selected the Batchawana Watershed due to the presence of the Turkey Lakes Watershed (TLW) experimental study^84^, which was established in 1979 and is one of the longest running watershed-based ecosystem studies in Canada^85^ The extensive scientific and support infrastructure at TLW hosts a comprehensive environmental data record which we utilized to estimate C fluxes and understand likely baseline conditions for our sites. The forests of the regions are composed of uneven-aged hardwood forests with primarily podzolic (spodosols) soils with well-developed forest floor horizons^86^. We focus this study on hardwood-dominated forests in the hemi-boreal region because of the expected shift towards hardwood species in temperate forests and the associated increased pressure on hardwood forests^87^, and because this forest region is expected to undergo particularly rapid climate shifts^88^. The most recent operational logging occurred in the 1950s when the area was selectively logged for high-quality yellow birch (*Betula alleghaniensis*), sugar maple (*Acer saccharum*), white spruce (*Picea glauca*), and white pine (*Pinus strobus*)^89^.

To select experimental catchments, the regional commercial forestry company (Boniferro Mill Works Inc., Sault Ste. Marie, Ontario) provided historical and future harvest plans. Within the future harvest region, we delineated catchments of headwater streams. Catchment boundaries were delineated using WhiteboxTools^90^, a python scripting API for geospatial analysis, along with 30m digital elevation models from the Government of Ontario and existing hydrological data from the Ontario Integrated Hydrology (OIH) dataset. Flow accumulation and flow pointer grids were generated using the D8 algorithm, where flow entering each cell is routed to only a single downstream neighbour^91^. Streams were extracted using the generated flow accumulation map and a threshold value of 0.128km^2^. This threshold value was selected by stepping through values and visually validating the generated network against satellite images of streams, catchments generated in smaller areas from 5m LiDAR-derived digital elevation models (DEMs) available for the TLW, and the stream network shape from the OIH dataset.

We then matched the harvest catchments to control catchments by maximising their similarity in a) topography, b) forest composition, and c) ecosite composition. Topography results were generated for each grid cell in the input DEM and included the slope gradient (i.e., steepness in degrees), aspect, sediment transport index, topographic wetness index and Pennock landform classification^101^. For each topographic variable we calculated median, maximum, minimum, and standard deviation values which were then assigned to the derived catchments. Forest composition was generated using the Ontario Forest Resources Inventory (FRI). This inventory is based on digital aerial photo interpretation and field surveys, with a recent incorporation of LiDAR data. From this layer we extracted polygons of overstory and understory species composition (and averaged the percent composition when both were available for a single site), tree species identified in the stand (or the uppermost canopy if the stand contained two or more distinct layers), and the percent cover each tree species occupies within the canopy. Additionally, we utilised the primary ecosite attribute from the FRI database. Ecosite is defined as an ecological unit comprised of relatively uniform geology, parent material, soils, topography, and hydrology and consists of related vegetation conditions. An intersection function was then used to calculate the percentage of area contributed by each tree species and ecosite to the catchment. We excluded possible control catchments based on a 300m distance to passable road (a semi-arbitrary threshold selected based on logistical concerns). Road data were downloaded from the Ministry of Natural Resources. Catchments within 50m of the DEM edge were also removed to avoid edge effects. These steps left n = 2177 catchments for further selection.

Control catchments were then selected using a divisive (top-down) hierarchical clustering analysis. This procedure is defined by a stepwise algorithm which starts with one cluster of all observations, and splits clusters into subgroups with each successive step based on within-group similarity and inter-group distance defined by the Ward’s method linkage and Euclidean distance, respectively. Permutations continued until there was only a single catchment remaining in the same branch/cluster as the catchments selected for harvest.

Within each of the four experimental catchments, hillslopes were further partitioned into four topographic features based on morphological features of surface DEMs according to Conacher and Dalrymple 1977^102^ and the Height Above the Nearest Drainage (HAND) terrain model^92^. The HAND model correlates with the depth of the water table, providing a spatial representation of soil water environments (k= 42 clusters)^93^. The four features were shoulder, backslope, footslope and toeslope.

### Logging

All harvest operations were performed according to the Ontario Stands and Site Guide for tolerant hardwood selection cuts^94^. Selection cut aims to remove, on average, 30% of the total basal area of tree stems in a stand but never more than 33%. Trees were felled by an Avery Tigercat LX380D feller buncher. Limbs were removed on site, and the tree-length stems were forwarded to roadside landings by rubber-tired skidders with tire chains. Operators were careful to avoid driving machinery into or across stream beds with adherence to the Ontario Stands and Site Guide^94^, including leaving 30m buffers on all mapped or obvious watercourses and avoidance of sensitive wet soils following Stands and Site Guide^89^.

### Stream water samples

Stream water samples were collected monthly during ice-free seasons beginning in September 2019. Surface water (500mL) was grab-sampled at the bottom of each hillslope at the channel head. DOM quantification and molecular characterization was performed on water filtered through 0.45 µm glass fibre syringe filters (Kinesis Inc, USA) into pre-combusted (4 h, 400 °C) amber vials and acidified to pH 2 with HCl within 24 hours. Samples were stored at 4 °C in the dark until analysis within four months. DOM was quantified using a Shimadzu TOC-L (Shimadzu, Japan) with concentrations determined via an L-arginine standard curve. For molecular characterization, we extracted 10 mL of each sample onto 100mg solid-phase extraction columns (Bond Elut PPL, Agilent) and eluted the sample with 3 mL methanol (ULC grade) using the methods described in Dittmar et al^95^. We also characterised DOM with optical analyses of filtered and refrigerated water, described above. Three-dimensional fluorescence scans were performed using a Cary Eclipse (Varian Instruments, USA), and absorbance measured with a Cary 60 UV–Vis (Agilent Technologies, USA). Fluorescence scans were run at 5 nm excitation steps from 250 to 450 nm, and emissions were read at 2 nm steps from 300 to 600 nm. Spectral corrections (instrument and inner-filter), and calculation of the humification index (HIX)^58^ were applied following standard procedures using the staRdom R package^96^. Finally, major ions, nutrients, and metal concentrations were measured from a subset of the sampled water at the Great Lakes Forestry Centre, Sault Ste. Marie, Ontario, according to methods outlined in ref^43^.

### Soil water collection

At all hillslope positions, at least 60 mL of soil water was sampled at 15, 30 and 60 cm depths with tension lysimeters. The lysimers consisted of 60-mm-long round bottom necked porous cups with an outer diameter of 48 mm and an effective pore size of 1.3 µm (model 0653X01-B02M2, Soilmoisture Equipment Corp). Lysimeters were installed 6 to 12 weeks prior to first sampling using a slurry of non-treated native soil. All lysimeters were evacuated at least twice prior to first sampling to allow for calibration^97^. The sampling bottles were evacuated to a negative pressure of 50 kPa with a hand pump so suction pressure was ca. 50 mbar above the actual soil water tension at collection. At 5 cm depths, lysimeters could not be securely installed. Therefore, we sampled pore water with micro-tensiometers designed to extract fluids non-destructively from soils using a vacuum^98^ through a 0.6 µm ceramic cup (Rhizon CSS samplers, Rhizosphere Research Products, Netherlands). All the water samplers were installed in triplicate and water pooled at each of the four depth and position combinations to retrieve sufficient volume for analysis. The hillside design of 16 soil samples and 1 stream sample was replicated across the 4 catchments for a total number of 68 samples per sampling date. Soil water samples were filtered through a 0.45 µm glass fibre filter, acidified (pH 2) and stored at 4°C prior to subsequent analyses of DOM concentration and molecular characterisation. Sample water for DOM fluorescence was refrigerated but not acidified.

### FT-ICR MS analysis

Fourier-transform ion cyclotron resonance mass spectrometry (FT-ICR MS) analysis was performed on a 15 T Solarix (Bruker Daltonics, USA). The system was equipped with an electrospray ionization source (ESI, Bruker Apollo II) applied in negative ionization mode. Samples were diluted to yield a concentration of ∼5 ppm in ultrapure water and methanol 50:50 (vol/vol). This dilution was filtered through pre-cleaned and rinsed 0.2 µm polycarbonate syringe filters before analysis performed in random order. Electrospray ionization in negative mode (Bruker Apollo II) was done at 200 °C and the capillary voltage was set to 4.5 kV. The sample was injected at a flow rate of 120 µL h^−1^, the accumulation time was set to 0.05 s, and 200 scans were co-added for each spectrum in a mass range of 92–2000 Da. Each spectrum was internally calibrated with lists of known masses. Mass spectra were exported from the Bruker Data Analysis software at a signal-to-noise ratio of 0 and molecular formulae were assigned using the ICBM Ocean tool^99^. The method detection limit was set to 3. Junction of mass lists along mass to charge ratios (m/z) was performed via fast join at a tolerance of 0.5 ppm while standard smooth and additional isotope tolerance was 10‰. Singlet peaks occurring only once in the dataset were removed, then molecular formulae were assigned with a tolerance of 0.5 ppm in the range m/z 0–1000 within the limits C_1–100_H_1–100_O_0–70_N_0–4_S_0–2_P_0–1_. Before statistical analysis, relative intensities were normalized to the sum of intensities per sample. Furthermore, molecular formulae were only retained in the dataset if they occurred at least three times across all samples.

For each molecular formulae, we calculated 10 traits related to molecular weight, stoichiometry, chemical structure, and oxidation state. These traits were molecular mass, the heteroatom class, double bond equivalents (DBE = number of rings plus double bonds to carbon, DBE^100^), carbon number (C), Standard Gibb’s Free Energy of carbon oxidation (GFE)^101^, nominal oxidation state of carbon (NOSC), O:C ratio, H:C ratio, and AI_mo_^102, 103^. Molecules were classified as CRAM where DBE:C was between 0.30-0.68, DBE:H was between 0.20-0.95 and DBE:O were between 0.77-1.75^104^. Formulae with AI_mod_ ≤ 0.5, 0.5< to ≤0.66, and >0.66 were defined as highly unsaturated and phenolics, polyphenolic and condensed aromatic respectively. Formulae with 1.5 ≤ H:C ≤ 2.0, O:C ≤ 0.9 and N = 0 were defined as aliphatic^105^. Formulae with O/C > 0.9 were defined as sugar-like whilst peptide-like were defined as 1.5 ≤ H/C ≤ 2.0, and N > 0^105^.

To compare DOM between each soil sample and stream at each time point, we calculated the Raup-Crick dissimilarity metric β_RC_ . This metric estimates whether pairwise samples are more different in composition than expected by chance, given random draws from the compound pool at a site. For any given pair of soil and stream samples, β_RC_ was calculated by comparing the observed number of shared molecular formulae to the number expected by randomly sampling the same number in each site from the entire compound pool following the protocol described in ref^99^. The probability of sampling a compound was based on its abundance across all soil sites. We repeated this sampling 1000 times to measure the similarity between the molecular composition of DOM in streams and soils compared to a null expectation.

### DOM yield and wood carbon estimates

Monthly DOM yield estimates were estimated for the logged experimental catchments by using monthly precipitation and runoff values (mm month^-1^) in the nearby TLW (Figure S3), with similar size (ranging from 3.5-43.0 ha, median = 2.14 ha, mean= 6.25 ha) and forest composition as our study catchments. Daily precipitation data were collected at 1.5 km south-east of the TLW boundary according to methods described in ref^107^ and averaged by month. Streamflow from the TLW catchments was estimated using stage-discharge relationships at 90° V-notch weirs, with stage measured at 10 minute intervals using water level recorders^107^. Stream stage was converted to mean daily discharge (L s^-1^) by dividing catchment area and computing the total runoff for each month, as described in ref^108^. Approximate annual DOM yields were estimated by multiplying the estimated monthly DOM by the estimated annual water yields.

We estimated the amount of tree carbon removed from each catchment using an estimate of aboveground phytomass for the region is described within ref^109^ Phytomass was then converted to carbon using a 50% (w/w) dry wood conversion based on the canopy dominant sugar maple^110^.

### Statistical analyses

We tested if stream water chemistry varied with logging. We fitted linear models to stream water chemistry (including chemodiversity) and stream water DOM. The time period (before or after logging) and treatment (control or impact, i.e., logging) were fixed effects and we included the interaction between these two variables. We also included the number of days since logging as a predictor as well as the three-way interaction, i.e., before-after × control-impact × time, as described in ref^111^. We accounted for repeat sampling of the same sites through time by including a site-level random effect and estimated a continuous autoregressive correlation structure to account for temporal autocorrelation within year and site. Models were fitted with the lme function in the nlme^112^ package in R^113^. When the predicted variable was a count (i.e., number of molecular formulae), we modelled the error terms using a Poisson distribution and estimated statistical models using the R package glmmTMB^114^. The Poisson error structure was not possible to include with nlme, but, concurrently, we did not use glmmTMB for all statistical models since it does not allow for a continuous correlation structure with time. In the case of glmmTMB, we instead used ordinal values (order of sample points in time) to model the autoregressive temporal correlation structure as properly modelling the error distribution was more important.

## Acknowledgements

We are thankful to K. Klapproth and I Ulber with FT-ICR-MS measurements. To Irena F. Creed for sending much-needed field support. Thank you to Mike Thompson and Jason McLellan for contributing the harvest in “harvest experiment”. We also thank the staff at the Natural Resources Canada – Canadian Forest Service (NRCan-CFS) for considerable support, including with logistics and advice (Paul Hazlett, Dan McKenney, John Pedlar, Jason Leach, Rob Fleming) and general field and laboratory assistance (J Schadenberg, D Chartrand, Caroline Emilson, Ken McIlwrick and many others). This work was funded by a Gates Cambridge Scholarship (OPP1144) awarded to E.C.F, a H2020 ERC Grant sEEIngDOM (804673) to A.J.T, and NRCan-CFS Cumulative Effects program awards to E.J.S.E.

## Notes

### Competing Interest Statement

The authors have declared no competing interest.

