## Supplement for "Logging disrupts the ecology of molecules in headwater streams"

**Supplementary Material**

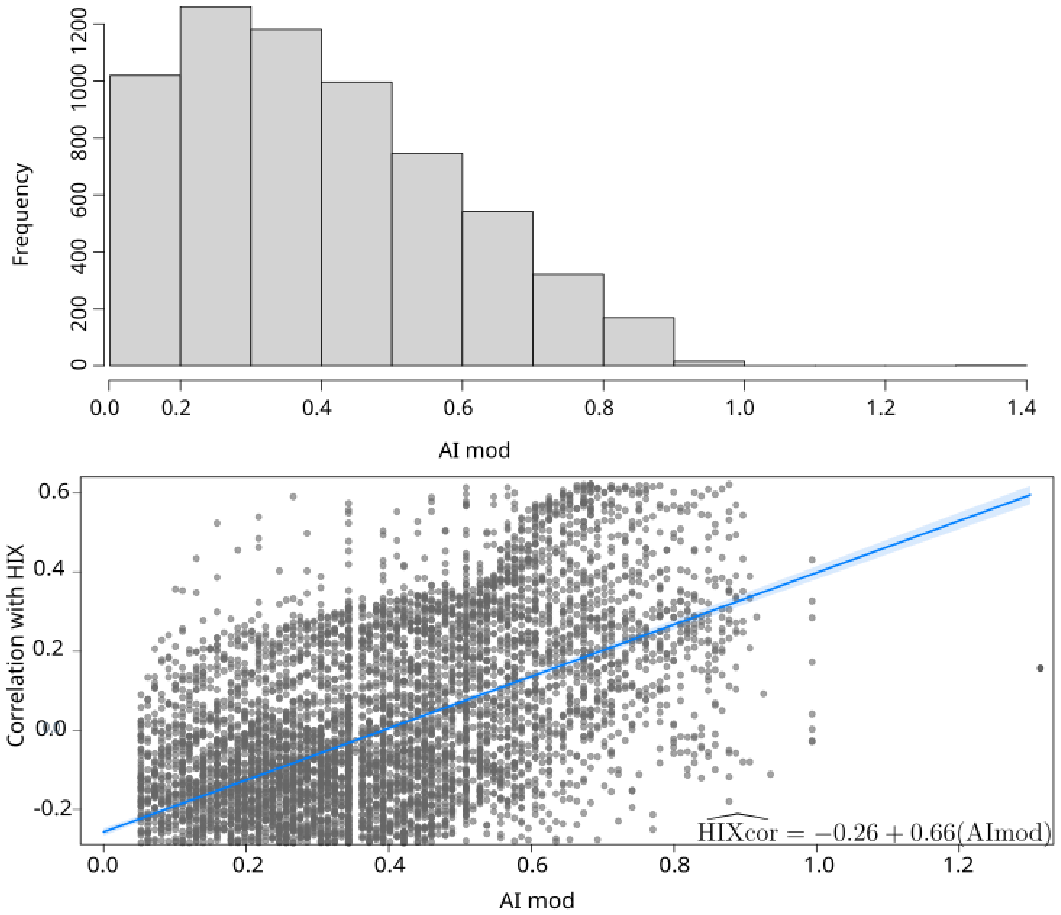

**Figure S1: Most compounds are likely bioavailable based on their modified aromaticity** ($AI_{mod}$)**.** $AI_{mod}$ in all molecular formulae (top) and their correlation with the humification index (HIX) (bottom). The reactivity of aromatic compounds (H:C< 1.1) is on average higher than reactivity in the "highly unsaturated" region (1.1 < H:C < 1.5)[1] and the uptake rate of DOM first decreases with increasing $AI_{mod}$, but at $AI_{mod}$ between 0.25 and 0.33 began to increase again.

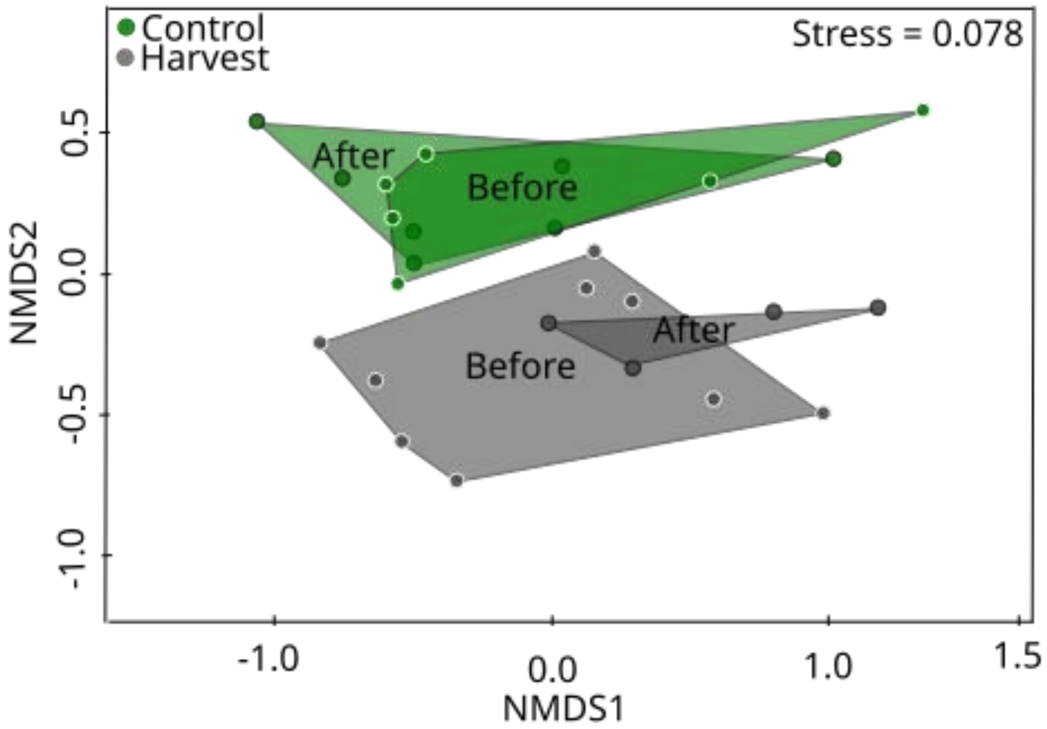

**Figure S2:** Non-metric multidimensional scaling (NMDS) ordination of DOM composition in streams before (June-September 2020, n=9) and after (October-November 2020, n=4) logging in replicate (n=4) catchments. The NMDS was fitted by calculating the Jaccard distance between observations from the presence-absence of molecular formulae. Compositional differences among groups were identified using a permutational multivariate analysis of variance ($R^2$ = 0.213 p = 0.029). Polygons were created by connecting the outermost data points within a grouping.

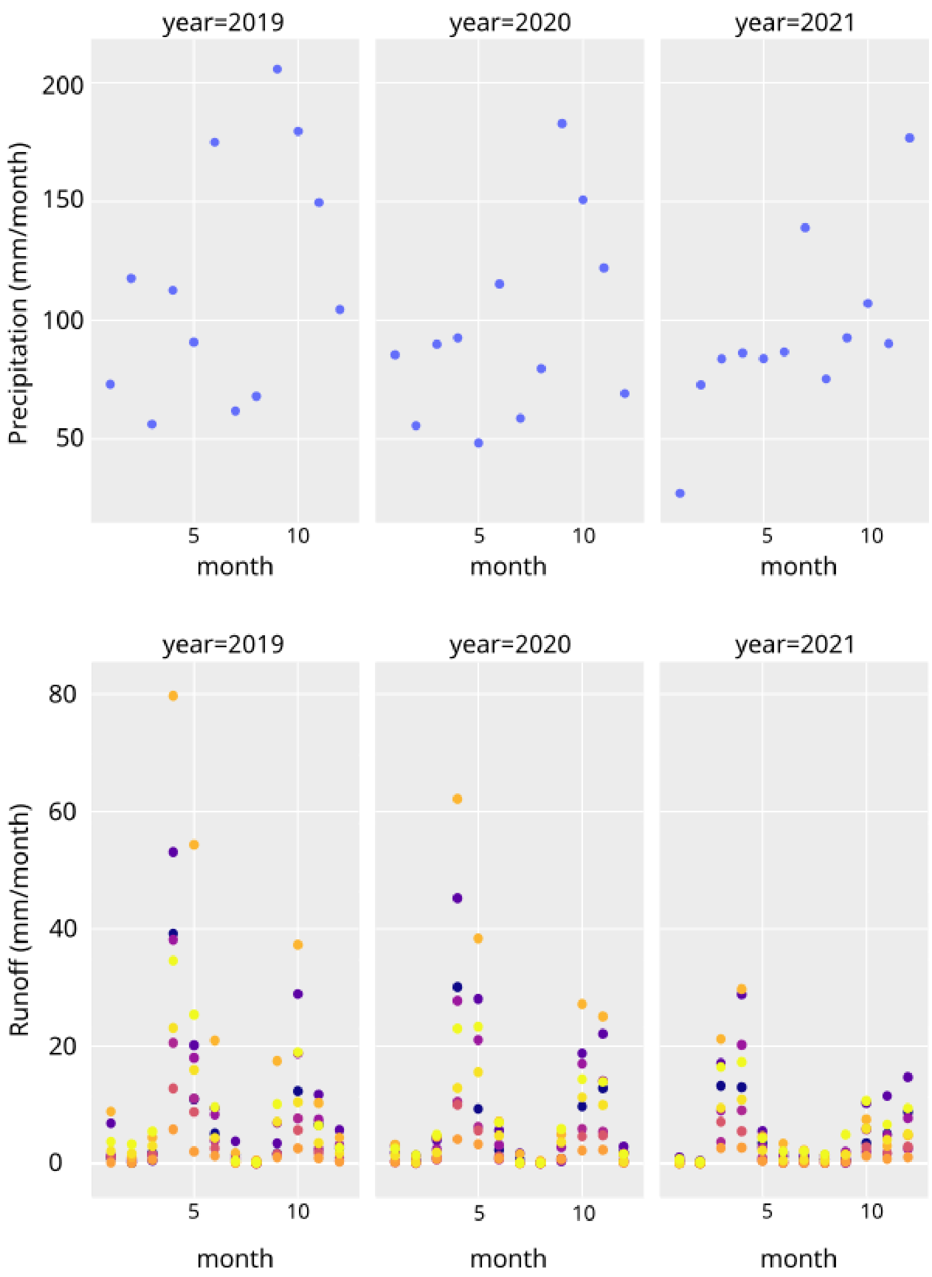

**Figure S3: Precipitation and runoff in the Turkey Lakes Watershed for each year.** Runoff was measured in each of 9 catchments (individual colours).

**Table S1: Estimated effects for models predicting the dissolved organic matter concentration in streams in the harvest year 2020).** Values in cells are the mean estimated effect relative to the intercept, with the intercept expressed relative to zero. Bolded effects do not overlap zero. CI are the 95% confidence interval. Model predictors were time (in days), Period (Before or After), and Treatment (Control or Harvest). Site was included as a random effect. σ is the within-site variance and τ is the between-site-variance. R-squared values are provided as marginal (considering only fixed) and conditional R-squared statistics (considering both fixed and random effects)[2].

| Fixed effects | Estimates | CI |
|---|---|---|
| Intercept (Before, Control) | **6.54** | 5.32 – 7.77 |
| Time (days) | **0.02** | 0.01 – 0.03 |
| After | -0.25 | -1.44 – 0.95 |
| Treatment | **-5.28** | -8.56 – -2.01 |
| Time × Before/After | **-0.07** | -0.11 – -0.03 |
| Time × Control/Harvest | **-0.02** | -0.04 – -0.00 |
| Before/After × Control/Harvest | 0.69 | -0.90 – 2.29 |
| Time × Before/After × Control/Harvest | **0.08** | 0.01 – 0.14 |
| Random Effect Estimates | | |
| σ | 0.41 | |
| τ | 0.08 | |
| N Sites | 4 | |
| Observations | 27 | |
| Marginal $R^2$ / Conditional $R^2$ | 0.86 / 0.91 | |

**Table S2: Estimated effects for models predicting the chemistry and fluorescence variables in streams in the harvest year (2020).** Model predictors were time (in days), period (Before or After), and treatment (Harvest or Control). Values in cells are mean estimated effects (standard error) relative to the intercept in parentheses, with the intercept expressed relative to zero. Bolded values were statistically significant at ***p <0.001, **p < 0.01, *p < 0.05. Site was included as a random effect to control for catchment-specific differences, σ is the within-site variance and τ is the between-site-variance. R-squared values are conditional R-squared statistics (taking both fixed and random effects into account)[126]. We also tested the fluorescence index (FI)[3], biological index (BIX)[4], quotient of molar absorptivity at 280 nm (E2E3)[5], and spectral slopes at 275–295 nm (S275–295), 350–350 nm (S350–400) and the spectral slope ratio (SR), absorbance at 254 and 300 (a254, a300), and concentrations of: total organic carbon (TOC), dissolved inorganic carbon, potassium, aluminum, iron, zinc, cadmium, nickel, and copper. None of the estimated effects in these models were significant. All additional absorbance measures were measured with the same methods as HIX (see Methods).

| | Fixed Effects | | | | | | Random Effects | | |
|---|---|---|---|---|---|---|---|---|---|
| Response | Intercept | Time | BA | CI | BA × CI | Time × BA × CI | σ | τ | $R^2$ |
| $NO_3$ | -0.01 (0.09) | **-4.57** (1.88) | 0.02 (0.10) | **0.57** (0.11) | -0.15 (0.13) | -2.28 (4.89) | 0.01 | 0.75 | 0.92/ 0.93 |
| $NH_4$ | -0.00 (0.01) | **-0.34** (0.13) | 0.01 (0.01) | 0.01 (0.01) | -0.01 (0.01) | -0.41 (0.34) | 0.03 | 0.05 | 0.34/ 0.39 |
| K | 0.09 (0.14) | **-8.27** (2.65) | **0.41** (0.13) | 0.37 (0.17) | **-0.47** (0.16) | -2.09 (6.60) | 0.02 | 0.75 | 0.42/ 0.42 |
| Cl | 0.04 (0.04) | -1.52 (0.75) | **0.13** (0.04) | 0.07 (0.04) | -0.10 (0.05) | -0.23 (1.95) | 0.06 | 0.09 | 0.39/ 0.45 |
| Al | 0.07 (0.04) | -0.79 (0.72) | **0.10** (0.04) | -0.06 (0.04) | -0.08 (0.04) | 0.30 (1.83) | 0.09 | 0.12 | 0.78/ 0.79 |
| Fe | 0.01 (0.04) | -1.24 (0.65) | **0.08** (0.03) | -0.00 (0.05) | **-0.08** (0.03) | -0.63 (1.53) | 0.07 | 0.22 | 0.37/ 0.40 |
| Zn | 11.03 (4.99) | 126.03 (99.62) | -3.07 (4.89) | -7.80 (6.12) | 1.26 (6.13) | **660.22** (251.46) | 20.45 | 5.53 | 0.27/ 0.32 |
| Cd | 0.14 (0.09) | -1.55 (1.88) | -0.19 (0.10) | -0.00 (0.11) | 0.11 (0.13) | **-14.10** (4.87) | 0.01 | 0.09 | 0.62/ 0.65 |
| Ni | 0.04 (0.16) | **-7.85** (3.29) | 0.08 (0.18) | 0.13 (0.19) | -0.13 (0.23) | -8.66 (8.52) | 0.02 | 0.75 | 0.44/ 0.49 |
| HIX | **18.71** (2.15) | 34.53 (25.78) | -5.75 (2.76) | **-13.54** (2.73) | **8.98** (3.83) | -106.73 (157.37) | 7.14 | 0.11 | 0.62/ 0.79 |

**Table S3: Estimated effects for models predicting the chemistry and fluorescence variables in streams between 2019-2021.** Model predictors were time (in days), Period (Before or After), and Treatment (Harvest or Control). Values in cells are mean estimated effects ± standard error relative to the intercept in parentheses, with the intercept expressed relative to zero. Bolded values were statistically significant at $p < 0.05$. Site and Year were selected as random effects to control for catchment-specific differences within year. σ is the within-site variance, τ1 is the between-group-variance by year, and τ2 is the between-group-variance by site. R-squared values are marginal R-squared statistics (taking only fixed effects into account). We also tested the fluorescence index (FI)[3], biological index (BIX)[4], quotient of molar absorptivity at 280 nm (E2E3)[5], and spectral slopes at 275–295 nm (S275–295), 350–350 nm (S350–400) and the spectral slope ratio (SR), absorbance at 254 and 300 (a254, a300), and concentrations of: total organic carbon (TOC), dissolved inorganic carbon, potassium, aluminum, iron, zinc, cadmium, nickel, and copper. None of the estimated effects in these models were significant. All additional absorbance measures were measured with the same methods as HIX (see Methods).

| | Fixed Effects | | | | | | Random Effects | | | |
|---|---|---|---|---|---|---|---|---|---|---|
| Response | Intercept | time | BA | CI | BA × CI | Time × BA × CI | σ | τ 1 | τ 2 | $R^2$ |
| $NO_3$ | **0.18** (0.09) | 0.40 (0.37) | **-0.13** (0.06) | 0.48 (0.12) | -0.15 (0.09) | 1.19 (0.77) | 0.01 | 0.00 | 0.10 | 0.93 |
| pH | **6.88** (0.17) | 0.07 (0.39) | **-0.25** (0.12) | 0.02 (0.23) | 0.02 (0.17) | **1.48** (0.67) | 0.03 | 0.19 | 0.04 | 0.49 |
| Conductivity | **37.08** (7.71) | **40.16** (11.33) | **-14.91** (3.46) | 16.69 (10.82) | -3.43 (4.80) | 28.69 (20.21) | 23.84 | 10.12 | 0.87 | 0.85 |
| Alkalinity | **0.25** (0.07) | **0.31** (0.14) | **-0.09** (0.04) | 0.15 (0.10) | -0.05 (0.06) | 0.13 (0.25) | 0.92 | 0.09 | 0.78 | 0.71 |
| Ca | **4.74** (0.96) | **4.10** (1.69) | **-1.90** (0.49) | 1.79 (1.34) | -0.45 (0.69) | 2.17 (3.12) | 0.49 | 1.21 | 0.20 | 0.76 |
| Mg | **0.79** (0.26) | **0.72** (0.34) | **-0.26** (0.10) | 0.52 (0.37) | -0.24 (0.14) | 0.60 (0.62) | 0.02 | 0.35 | 0.04 | 0.81 |
| Na | **1.04** (0.26) | 0.60 (0.34) | **-0.25** (0.11) | 0.73 (0.37) | **-0.31** (0.15) | 0.34 (0.59) | 0.02 | 0.35 | 0.38 | 0.85 |
| $SO_4$ | **2.88** (0.45) | 2.46 (1.23) | **-1.23** (0.35) | 0.14 (0.62) | 0.84 (0.48) | 1.49 (2.31) | 0.22 | 0.48 | 0.11 | 0.54 |

| | | | | | | | | | | |
|---|---|---|---|---|---|---|---|---|---|---|
| Mn | **11.40** (2.90) | **24.88** (11.76) | -5.76 (3.57) | -8.15 (3.88) | 5.83 (4.88) | 9.35 (21.04) | 23.16 | 1.11 | 0.10 | 0.27 |
| Pb | **0.13** (0.03) | 0.00 (0.14) | **-0.10** (0.04) | -0.05 (0.04) | 0.04 (0.06) | -0.20 (0.25) | 0.87 | 0.17 | 0.80 | 0.38 |
| HIX | **18.37** (2.22) | 34.10 (24.17) | **-5.54** (2.27) | **-14.24** (2.72) | **8.57** (2.78) | 28.16 (26.82) | 6.27 | 0.00 | 1.36 | 0.78 |

**Table S4: Compound counts in treatments.** All values were estimated by mixed effects models. CI is the 95% confidence interval.

| Treatment | Estimates | CI |
| --- | --- | --- |
| Before, Control | **5386.86** | 4733.01 – 6131.57 |
| Before, Harvest | **4771.52** | 3999.31 – 5692.08 |
| Control, After | **4152.74** | 364095 – 4738.82 |
| Harvest, After | **6292.24** | 5521.74 – 7172.08 |
| Model statistics | | |
| N Site | 4 | |
| Observations | 24 | |
| Marginal $R^2$ | 0.99 | |

**Table S5: Pairwise dissimilarity between matched control sites and harvests before and after logging treatment.** All values were estimated by a linear model with site as a fixed effect to control for site-level differences. Bolded effects do not overlap zero. CI is the 95% confidence interval.

| Predictors | Estimates | CI |
| --- | --- | --- |
| Before | **0.11** | 0.02 – 0.19 |
| After | **0.12** | 0.00 – 0.24 |
| $R^2$ / $R^2$ adjusted | 0.21 / 0.16 | |

**Table S6: Percentages of each elemental class of the total number of compounds in each group of compounds.** The total number of molecules is given in parentheses. Elemental classes are not mutually exclusive; therefore, rows can sum to >100. We separately classified compounds that appeared in the harvest site only after logging (harvest gains); compounds that were absent from the harvest site after logging (harvest losses); all compounds present in all streams (all) and all compounds present in the harvest site before logging.

| Group (n) | N | S | P | NS | NP |
|---|---|---|---|---|---|
| Harvest Gains (1035) | 42.00 | 9.18 | 3.67 | 0.29 | 0.19 |
| Harvest Losses (320) | 24.38 | 21.88 | 19.06 | 0.31 | 0.31 |
| All (7444) | 32.28 | 10.45 | 7.27 | 0.63 | 0.09 |
| Before Harvest (4927) | 28.76 | 7.53 | 7.54 | 0.09 | 0.06 |

**Table S7: Percentages of each compound class of the total number of compounds in each group of compounds.** The total number of molecules is given in parentheses. We separately classified compounds into groups that appeared in the harvest site only after logging (harvest gains) and all compounds present in the harvest site before logging.

| Group (n) | CHO | CHOP | CHOS | CHONS | CHOSP | CHONP | CHON |
|---|---|---|---|---|---|---|---|
| Harvest Gains (1035) | 45.12 | 3.03 | 8.40 | 0.20 | 0.49 | 0.00 | 42.77 |
| Harvest Losses (320) | 36.33 | 24.1 | 20.26 | 18.65 | 0.00 | 0.32 | 0.32 |
| All (7444) | 51.39 | 6.63 | 9.01 | 0.33 | 0.57 | 0.04 | 32.02 |
| Before Harvest (4927) | 54.00 | 7.36 | 6.99 | 0.06 | 0.71 | 0.06 | 30.82 |

**Table S8 The difference in mean soil-stream similarity before and after harvest in treatment and control sites.** Estimated mean (± 95% CI) pairwise similarity between soil and stream sites before (intercept) and after logging treatment in each of the control and harvest. CI are the 95% confidence interval.

| **Harvest** | **Similarity (Soil-Stream)** | |
|---|---|---|
| Predictors | *Estimates* | *CI* |
| Before Harvest | **0.61** | 0.55 – 0.67 |
| After Harvest | -0.04 | -0.11 – 0.03 |
| Observations | 41 | |
| **Control** | ***Similarity (Soil-Stream)*** | |
| Before Harvest | **0.60** | 0.58 – 0.62 |
| After Harvest | **-0.07** | -0.10 – -0.04 |
| Observations | 42 | |

**Table S9: The difference in soil (depth x position)-stream similarity before and after harvest in the treatment and controls.** Estimated mean (± 95% CI) pairwise similarity between soil and stream sites after logging treatment (compared to before) in each of the control and harvest, at each soil position (depth × position**)**. CI are the 95% confidence interval. Bolded effects do not overlap zero.

| Predictors | Estimates | CI |
|---|---|---|
| Shoulder, 5 cm | -0.08 | -0.79-0.11 |
| **Shoulder, 15cm** | **0.20** | **2.23-0.09** |
| Shoulder, 30 cm | -0.02 | -0.23-0.12 |
| Shoulder, 60cm | -0.03 | -0.22-0.13 |
| Backslope, 5 cm | 0.09 | 0.09-0.95 |
| Backslope, 15cm | -0.09 | -0.57-0.17 |
| Backslope, 30cm | -0.02 | -0.21-0.07 |
| **Backslope, 60cm** | **0.13** | **0.57-2.31** |
| Footslope, 5cm | 0.08 | 0.09-0.91 |
| **Footslope, 15cm** | **0.25** | **0.09-2.74** |
| Footslope, 30cm | 0.03 | 0.13-0.27 |
| Footslope, 60cm | 0.15 | 0.13-1.18 |

| | | |
|---|---|---|
| Toeslope, 5cm | 0.15 | 0.14-1.12 |
| Toeslope, 15cm | 0.11 | 0.09-1.19 |
| Toeslope, 30cm | 0.09 | 0.08-1.20 |
| Toeslope, 60cm | 0.03 | 0.09-0.36 |
| Observations | 4 | |
